# Fusion-oncogenes are associated with increased metastatic capacity and persistent disease in pediatric thyroid cancers

**DOI:** 10.1101/2021.07.23.453235

**Authors:** Aime T. Franco, Julio C. Ricarte-Filho, Amber Isaza, Zachary Jones, Neil Jain, Sogol Mostoufi-Moab, Lea Surrey, Theodore W. Laetsch, Marilyn M. Li, Jessica Clague DeHart, Erin Reichenberger, Deanne Taylor, Ken Kazahaya, N. Scott Adzick, Andrew J. Bauer

**Affiliations:** Division of Endocrinology and Diabetes, Children’s Hospital of Philadelphia, Philadelphia, PA; Division of Oncology, Children’s Hospital of Philadelphia, Philadelphia, PA; Department of Pathology and Laboratory Medicine, Children’s Hospital of Philadelphia, Philadelphia, PA; School of Community and Global Health, Claremont Graduate University, Claremont, CA; Department of Biomedical and Health Informatics, Children’s Hospital of Philadelphia, Philadelphia, PA; Division of Pediatric Otolaryngology, Children’s Hospital of Philadelphia, Philadelphia, PA; Department of Surgery, Children’s Hospital of Philadelphia, Philadelphia, PA

## Abstract

**Background:** In 2014, data from a comprehensive multiplatform analysis of 496 adult papillary thyroid cancer samples reported by The Cancer Genome Atlas project suggested that reclassification of thyroid cancer into molecular subtypes, *RAS*-like and *BRAF*-like, better reflects clinical behavior than sole reliance on pathological classification. The aim of this study was to categorize the common oncogenic variants in pediatric differentiated thyroid cancer and investigate if mutation subtype classification correlated with the risk of metastasis and response to initial therapy in pediatric DTC.

**Methods:** Somatic cancer gene panel analysis was completed on DTC from 131 pediatric patients. DTC were categorized into *RAS*-mutant (*H-K-NRAS*), *BRAF*-mutant (BRAF p.V600E) and *RET/NTRK* fusion (*RET, NTRK1* and *NTRK3* fusions) to determine differences between subtype classification in regard to pathological data (AJCC TNM) as well as response to therapy 1-year after initial treatment had been completed.

**Results:** Mutation-based subtype categories were significant in most variables, including age at diagnosis, metastatic behavior, and the likelihood of remission at 1-year. Patients with *RET/NTRK* fusions were significantly more likely to have advanced lymph node and distant metastasis and less likely to achieve remission at one year than patients within *RAS*- or *BRAF-mut* subgroups.

**Conclusions:** Our data supports that genetic subtyping of pediatric DTC more accurately reflects clinical behavior than sole reliance on pathological classification with patients with *RET/NTRK* fusions having worse outcomes than those with *BRAF*-mutant disease. Future trials should consider inclusion of molecular subtype into risk stratification.

## BACKGROUND

In 2014, the Thyroid Cancer Genome Atlas (TCGA) reported that classification of adult PTC into molecular subtypes based on a mRNA expression signature, *RAS*-like and *BRAF*-like, more accurately reflected cellular signaling, cellular differentiation and clinical behavior when compared to histology alone.^1^ This observation has fueled debate whether identification of oncogenic alterations more accurately predict the risk of malignancy in thyroid nodules with indeterminate cytology may be used to stratify therapy, including the extent of surgery, lobectomy versus total thyroidectomy, as well as central compartment lymph node dissection.^2,3^

With the reduced costs of next-generation sequencing, the use of somatic cancer gene panel analysis in pediatric patients with differentiated thyroid cancer (DTC) has expanded with current data showing a shifted distribution of driver alterations with a higher incidence of oncogenic fusions rather than point mutations in children and adolescents compared to adults.^4,5^ Similar to the TCGA based molecular sub-type classification system based on mRNA expression signatures, we sought to determine whether oncogenic subtyping based on identified mutations or fusions predicts phenotypic behavior and outcomes in pediatric patients with DTC.

## METHODS

### Patient population

Our series comprised 131 pediatric thyroid tumors (122 PTCs and 9 FTCs) from surgical specimens. This included 66 surgical specimens sequentially collected from 2016-2019 and analyzed in the Department of Pathology at Children’s Hospital of Philadelphia (CHOP) as well as 65 archived surgical specimens collected from 1989-2012 that were previously genotyped on a commercial platform.^6^ All of the tumors were sequentially collected and analyzed without bias selection limited only by sample adequacy. The research protocol was approved by CHOP’s Institutional Review Board.

DTC classification was based on standard histopathological criteria defined by the World Health Organization.^7^ Tumors were staged according to the 7^th^ edition of the American Joint Committee on Cancer (AJCC) staging manual.^8^ Invasion was defined as spread to regional lymph nodes and/or distant metastasis. Remission was assessed at one year after surgery and defined as a basal-thyroglobulin (Tg) below the lower limit of detection and no evidence of persistent thyroid cancer on radiological or radioiodine whole-body scan (RAI-WBS). Neck ultrasound was used for patients with disease limited to the neck and undetectable Tg. Chest CT was added for patients with a history of pulmonary metastasis on initial imaging. RAI-WBS was used to assess for persistent disease in patients with detectable Tg (> 10 ng/ml and/or increasing trend) and no evidence of persistent thyroid cancer based on neck US and chest CT.

### Sequencing platform and variant calling

The 66 tumors collected between 2016 and 2019 were sequenced as part of routine clinical care using both the CHOP Solid Tumor Panel (CSTP) and CHOP Cancer Fusion Panel (CCFP). CSTP is a targeted next-generation sequencing (NGS) assay encompassing 238 cancer genes. The assay is designed to detect single nucleotide variants (SNVs), indels, and copy number alterations (CNAs) as described previously.^9^ Briefly, genomic DNA was extracted from the tumor samples and libraries were prepared using probes targeting 238 genes and sequenced on HiSeq platform using 150 bp paired-end sequencing. Sequence data were analyzed using the home brew software ConcordS v2 (for SNVs and indels) and NextGENe v2 NGS Analysis Software (for CNAs; SoftGenetics, LLC, State College, PA). Fusion gene detection was performed using the CHOP Cancer Fusion Panel as previously described.^10^ Briefly, target-specific primers covering 673 exons were custom designed to identify known fusion genes and potential novel fusion genes associated with 110 cancer genes using Anchored Multiplex PCR (AMP™) technology (ArcherDX, Inc. Boulder, CO). The 65 cases collected between 1989 and 2012 had been previously genotyped using Asuragen’s first-generation thyroid test, miRinform^®^ Thyroid Test, as described.^6^ This panel interrogates the presence of the most common mutations in *BRAF*, *HRAS*, *KRAS* and *NRAS* and three fusion transcripts (*RET/PTC1, RET/PTC3* and *PAX8/PPARG*). The miRinform^®^ panel did not include analysis for *DICER1* mutations or NTRK-fusions. Unfortunately, tissue from the miRinform^®^ cohort was not available for repeat analysis using the more comprehensive CHOP panels. All cancer genes included in the CSTP and CCFP panels and the miRInform Thyroid Test are listed in **Supplementary Table 1**.

Mutations were subcategorized into three groups, *RAS*-mutant (*H/K/NRAS* mutations and *PAX8/PPARG* fusions), *BRAF*-mutant (BRAF p.V600E mutations), or *RET/NTRK* fusions (*RET, NTRK1* and *NTRK3* fusions) based on previous published reports in adults^1,2,11^ as well as pediatric data showing genotype-associated differences in invasive behavior.^12–14^ Tumors with no identified genetic alteration in the miRInform Thyroid Test, or genetic alterations classified as not previously identified as driver mutations in pediatric DTC with the CSTP panel, were characterized as indeterminate.^9^

### Data analyses

Data analysis was performed using Statistical analyses were run in R 4.0.5 and R Studio 1.4.1106. Frequencies and proportions were used as descriptive statistics for categorical variables. Mutation status was the primary variable of interest and so this was explored over a number of different dimensions of the data. Associations between covariates and mutation status were tested using Fisher’s exact tests, to account for small cell sizes. A two-sided p value of less than 0.05 was considered statistically significant. Mutation data and clinicopathological characteristics from adult papillary thyroid cancer was collected from The Cancer Genome Atlas (TCGA) Data Portal (https://tcga-data.nci.nih.gov).

## RESULTS

### Patient Demographics and Thyroid Pathology

Clinicopathologic characteristics of the study population are summarized in **Table 1**. The study cohort included 131 samples from 97 (74%) female patients and 34 (26%) male patients with a mean age of 14.53 ± 2.99 years. Tumors were divided based on their histopathological features: 60 (45.8%) classic variant (cPTC), 29 (22.1%) follicular variant (fvPTC), 14 (10.7%) diffuse sclerosing variant (dsvPTC), 8 (6.1%) mixed cPTC and fvPTC, 11 (8.4%) were other forms of PTC, including cribriform-morular variant (cmvPTC), solid variant (svPTC), and Warthin-like (WLPTC), and 9 (6.9%) follicular thyroid carcinoma (FTC).

**Table 1.**
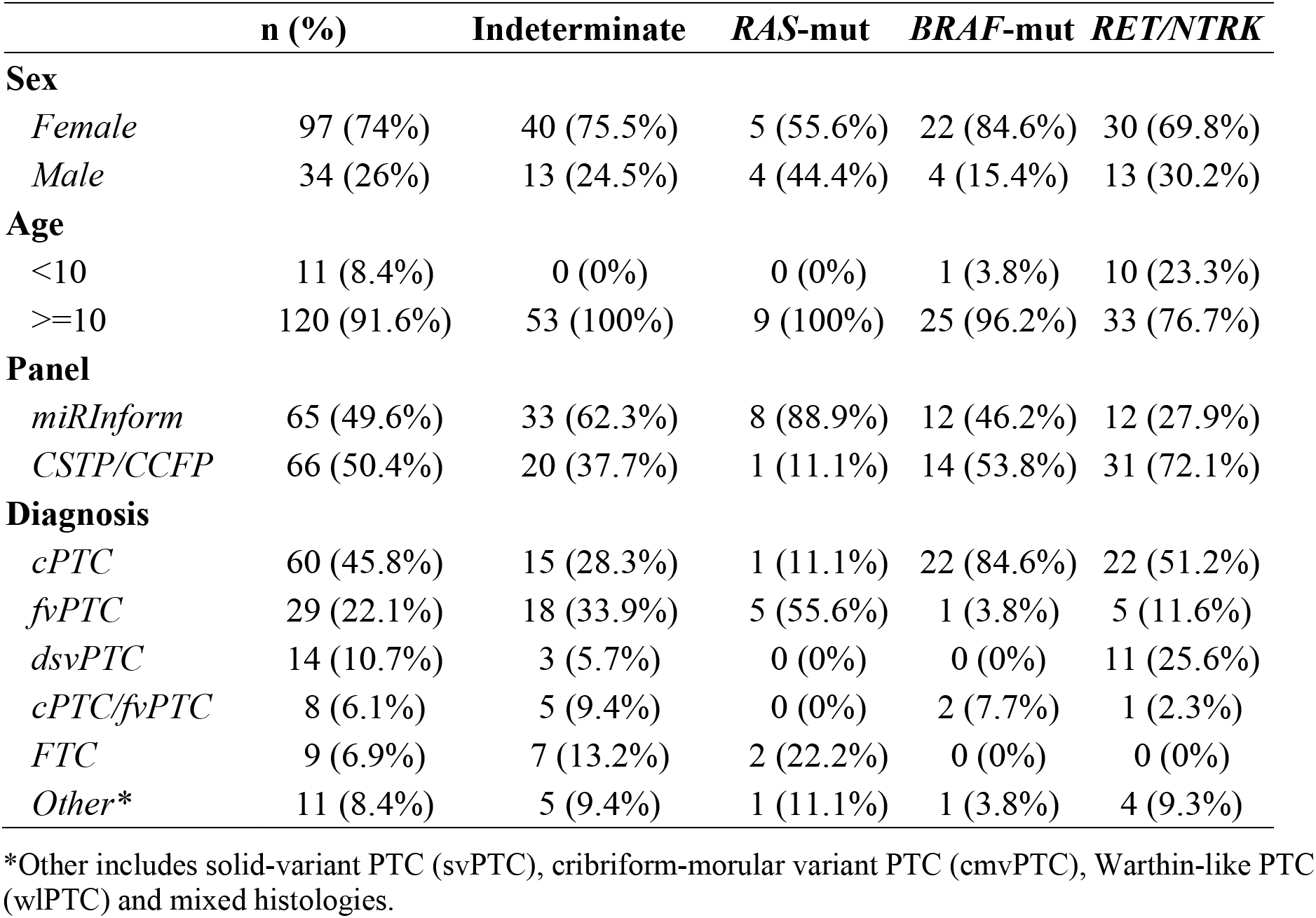
Demographics of 131 pediatric and adolescent patients.

|  | **n (%)** | **Indeterminate** | ***RAS*-mut** | ***BRAF*-mut** | ***RET/NTRK*** |
| --- | --- | --- | --- | --- | --- |
| **Sex** |  |  |  |  |  |
| *Female* | 97 (74%) | 40 (75.5%) | 5 (55.6%) | 22 (84.6%) | 30 (69.8%) |
| *Male* | 34 (26%) | 13 (24.5%) | 4 (44.4%) | 4 (15.4%) | 13 (30.2%) |
| **Age** |  |  |  |  |  |
| <10 | 11 (8.4%) | 0 (0%) | 0 (0%) | 1 (3.8%) | 10 (23.3%) |
| >=10 | 120 (91.6%) | 53 (100%) | 9 (100%) | 25 (96.2%) | 33 (76.7%) |
| **Panel** |  |  |  |  |  |
| *miRInform* | 65 (49.6%) | 33 (62.3%) | 8 (88.9%) | 12 (46.2%) | 12 (27.9%) |
| *CSTP/CCFP* | 66 (50.4%) | 20 (37.7%) | 1 (11.1%) | 14 (53.8%) | 31 (72.1%) |
| **Diagnosis** |  |  |  |  |  |
| *cPTC* | 60 (45.8%) | 15 (28.3%) | 1 (11.1%) | 22 (84.6%) | 22 (51.2%) |
| *fvPTC* | 29 (22.1%) | 18 (33.9%) | 5 (55.6%) | 1 (3.8%) | 5 (11.6%) |
| *dsvPTC* | 14 (10.7%) | 3 (5.7%) | 0 (0%) | 0 (0%) | 11 (25.6%) |
| *cPTC/fvPTC* | 8 (6.1%) | 5 (9.4%) | 0 (0%) | 2 (7.7%) | 1 (2.3%) |
| *FTC* | 9 (6.9%) | 7 (13.2%) | 2 (22.2%) | 0 (0%) | 0 (0%) |
| *Other\** | 11 (8.4%) | 5 (9.4%) | 1 (11.1%) | 1 (3.8%) | 4 (9.3%) |

\*Other includes solid-variant PTC (svPTC), cribriform-morular variant PTC (cmvPTC), Warthin-like PTC (wlPTC) and mixed histologies.

### Identification and grouping of genetic alterations

Of the 131 patient samples, genetic alterations were identified in 78 of the 131 tumors; 9 with a *RAS*-mutant (6.9%, 9/131), 26 with *BRAF* (19.8%, 26/131), and 43 with a *RET*- or *NTRK*-fusion (32.8%, 43/131; **Table 1**). There were 53 tumors classified as indeterminate. Secondary to more limited coverage of oncogenic driver alterations, there were more tumors classified as indeterminate in the miRInform group compared to the CSTP/CCFP group (62.3% vs 37.7%) most notably fewer *RET/NTRK*-fusions identified in the miRInform group compared to the CSTP/CCFP group (27.9% vs 72.1%; **Table 1**).

**Figure 1** summarizes the mutational landscape of pediatric DTC and clinicopathological features. As previously reported, we found a correlation between genotype and pathological phenotype.^12–14^ *RAS*-mutations and *PAX8-PPARG* fusions were more commonly associated with fvPTC and FTCs than the other genotypes. There were 4 NRAS p.Q61R and 1 *PAX8-PPARG* fusions in five encapsulated fvPTC, one NRAS p.Q61R in a cPTC, one *PAX8/PPARG* in a cPTC/fvPTC/svPTC mixed histology and a single HRAS p.Q61R and single KRAS p.G12V in two FTC samples. The BRAF p.V600E mutation was most commonly associated with cPTC, observed in 26 (19.8%) samples. *RET/NTRK* fusions were found in 43 (32.8%) tumor samples spread across various subtypes of PTC, including 22 cPTC samples, 11 dsvPTC, 5 fvPTC, 1 svPTC, 1 fvPTC/svPTC, 1 cPTC/fvPTC and 2 cPTC/fvPTC/svPTC. The fusions included 20 *RET/PTC1* (*CCDC6-RET*), 1 *RET/PTC2* (*PRKAR1A-RET*), 8 *RET/PTC3* (*NCOA4-RET*), 2 *SPECC1L-RET*, 1 *EML4-RET*, 1 *TRIM24-RET*, 1 *CCDC186-RET*, 2 *TPR-NTRK1*, 1 *IRF2BP-NTRK1*, 1 *SQSTM1-NTRK1* and 5 *ETV6-NTRK3*.

**Figure 1.**
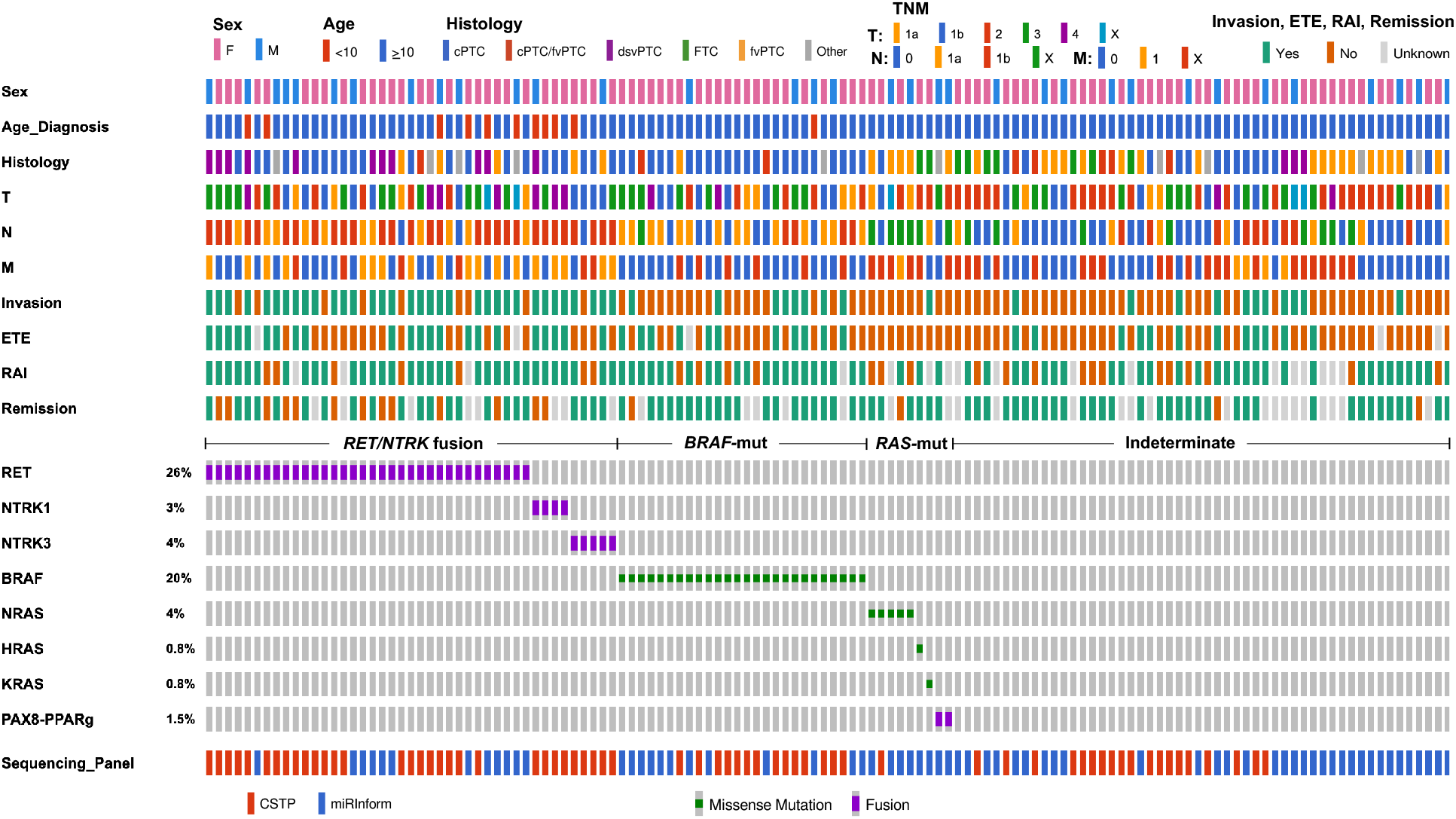
Genetic landscape and clinicopathological characteristics of 131 pediatric thyroid cancers. Clinicopathological characteristics includes sex, age, histology, tumor (T) status, lymph node (N) metastasis status, distant metastasis status (M), invasion, extrathyroidal extension (ETE), radioactive iodine (RAI) therapy and remission. The most frequent genetic alterations (fusion oncogenes and mutations) and their prevalence are shown. Genetic alterations were categorized in *RET/NTRK* fusions (*RET* and *NTRK1/3* fusions), *BRAF*-mut (BRAF p.V600E) and *RAS*-mut (*H-K-NRAS* and *PAX8/PPARG*).

Of note, 4 additional potential kinase activating in-frame fusions (2 *TFG-MET*, 1 *TG-FGFR1* and 1 *PRKD2-BRAF*) and 3 mutations (1 BRAF p.T599del and 2 TSHR p.M453T and p.D633Y) were identified by the more comprehensive CSTP/CCFP panel. In addition, we found mutations associated with increased risk of thyroid cancer: 3 cases harbored alterations of *APC* (3 cmvPTC) and 2 cases harbored biallelic mutations of *DICER1* (1 fvPTC and 1 FTC). The 12 cases above were classified as indeterminate due to their low prevalence and uncertain molecular category. All genetic drivers found in this study are listed in **Supplementary Table 2**.

### Relationship between genetic alterations and clinical characteristics

The genetic alterations and correlation with clinicopathologic characteristics and outcomes are summarized in **Table 2**. Several strong associations between the covariates and mutation categories were observed. When comparing the 4 categories of mutation status: indeterminate, *RAS*-mutant, *BRAF*-mutant, and *RET/NTRK* fusion, all variables were found to be statistically significant (p-value <0.05), except for sex and AJCC T (tumor size) category. There were very few *RAS*-mutant samples, therefore we restricted analysis and comparison to *BRAF*-mutant and *RET/NTRK* fusions. After restricting the sample to those with *BRAF*-mutant vs *RET/NTRK* fusion status, statistically significant associations were still found among age, AJCC N (lymph node metastasis) and M (distant metastasis, all pulmonary) categories as well as remission at 1-year (p-value <0.05). Significantly, no distant metastasis was detected in any patients with *BRAF*-mutant thyroid tumors. Thirty-seven percent of patients with *RET/NTRK* fusion tumors did not attain remission at one year compared to less than 5% of patients with a tumor harboring a *BRAF* mutation.

**Table 2.** Association between the covariates and mutations categories: *BRAF-mut, RAS-mut, RET/NTRK* fusion and Indeterminate.

|  |  | Indeterminate | *RAS*-mut | *BRAF*-mut | *RET/NTRK* | p-value,<br>4 Categories | p-value,<br>*BRAF* vs<br>*RET/NTRK* |
| --- | --- | --- | --- | --- | --- | --- | --- |
| ETE | Yes | 13 (24.5%) | 0 (0%) | 12 (46.2%) | 23 (53.5%) | **2.66E-03** | 8.03E-01 |
|  | No | 40 (75.5%) | 9 (100%) | 14 (53.8%) | 20 (46.5%) |  |  |
| Invasion | Yes | 11 (20.8%) | 1 (11.1%) | 8 (30.8%) | 34 (79.1%) | **2.13E-08** | **1.28E-04** |
|  | No | 42 (79.2%) | 8 (88.9%) | 18 (69.2%) | 9 (20.9%) |  |  |
| Histology | FTC | 7 (13.2%) | 2 (22.2%) | 0 (0%) | 0 (0%) | **5.37E-03** | 1.00E+00 |
|  | PTC | 46 (86.8%) | 7 (77.8%) | 26 (100%) | 43 (100%) |  |  |
| Remission | Yes | 35 (94.6%) | 6 (85.7%) | 20 (95.2%) | 22 (62.9%) | **8.74E-04** | **9.08E-03** |
|  | No | 2 (5.4%) | 1 (14.3%) | 1 (4.8%) | 13 (37.1%) |  |  |
| Sex | Female | 40 (75.5%) | 5 (55.6%) | 22 (84.6%) | 31 (72.1%) | 3.70E-01 | 3.75E-01 |
|  | Male | 13 (24.5%) | 4 (44.4%) | 4 (15.4%) | 12 (27.9%) |  |  |
| Age | <10 | 0 (0%) | 0 (0%) | 1 (3.8%) | 10 (23.3%) | **5.98E-03** | 3.75E-01 |
|  | ≥10 | 53 (100%) | 9 (100%) | 25 (96.2%) | 33 (76.7%) |  |  |
| M | 0 | 27 (50.9%) | 2 (22.2%) | 20 (76.9%) | 19 (44.2%) | **5.86E-06** | **1.92E-04** |
|  | 1 | 4 (7.5%) | 1 (11.1%) | 0 (0%) | 18 (41.9%) |  |  |
|  | X | 22 (41.5%) | 6 (66.7%) | 6 (23.1%) | 6 (13.9%) |  |  |
| N | 0 | 28 (52.8%) | 2 (22.2%) | 7 (26.9%) | 3 (7.0%) | **4.09E-09** | **1.03E-04** |
|  | 1a | 7 (13.2%) | 1 (11.1%) | 14 (53.8%) | 11 (25.6%) |  |  |
|  | 1b | 9 (17.0%) | 0 (0.0%) | 4 (15.4%) | 29 (67.4%) |  |  |
|  | X | 9 (17.0%) | 6 (66.7%) | 1 (3.8%) | 0 (0%) |  |  |
| T | 1a | 4 (7.5%) | 2 (22.2%) | 4 (15.4%) | 4 (9.3%) | 1.12E-01 | 8.70E-01 |
|  | 1b | 8 (15.1%) | 1 (11.1%) | 7 (26.9%) | 10 (23.3%) |  |  |
|  | 2 | 24 (45.3%) | 4 (44.4%) | 5 (19.2%) | 6 (14.0%) |  |  |
|  | 3 | 13 (24.5%) | 1 (11.1%) | 8 (30.8%) | 14 (32.6%) |  |  |
|  | 4 | 2 (3.8%) | 0 (0%) | 2 (7.7%) | 7 (16.3%) |  |  |
|  | X | 2 (3.8%) | 1 (11.1%) | 0 (0%) | 2 (4.7%) |  |  |

cPTC variant was the most common histology for both *BRAF*-mut (n=22) and *RET/NTRK* fusion subgroups (n=22; 16 *RET*, 3 *NTRK1*, 3 *NTRK3* fusions). Even within this histological variant, we observed significant differences in metastatic behavior and remission status between these molecular subgroups. Metastases to lateral neck lymph nodes were found in 4/22 (18.2%) of patients with a *BRAF* mutation and 13/22 (59.1%) of patients with *RET/NTRK* fusions. Distant metastasis was present in 8/22 (36.4%) of the *RET/NTRK* fusion subgroup and none of the patients within *BRAF*-mut subgroup. Moreover, persistent disease at 1-year was more frequent in the subgroup harboring *RET/NTRK* fusions (5/15; 33.3%) than those with mutations in *BRAF* (1/18; 5.6%) (**Supplementary Table 3**).

Consistent with previously published results, a greater percentage of pediatric patients presented with nodal and distant metastasis compared to adult tumors reported in the TCGA database (**Figure 2**).^15^ In adult DTC samples, lateral lymph node metastasis was slightly increased in patients with *RET/NTRK* fusions vs *BRAF*-mut, but the difference is less pronounced than in our pediatric population (**Figure 2**). By contrast, adults harboring *RAS*-mutant PTC rarely present with lymph node or distant metastasis.^16^ This also seems to be the case for pediatric patients with *RAS*-mutant DTC, although the significance of this is unknown due to the limited number of *RAS*-mutant DTCs in our cohort.

**Figure 2.**
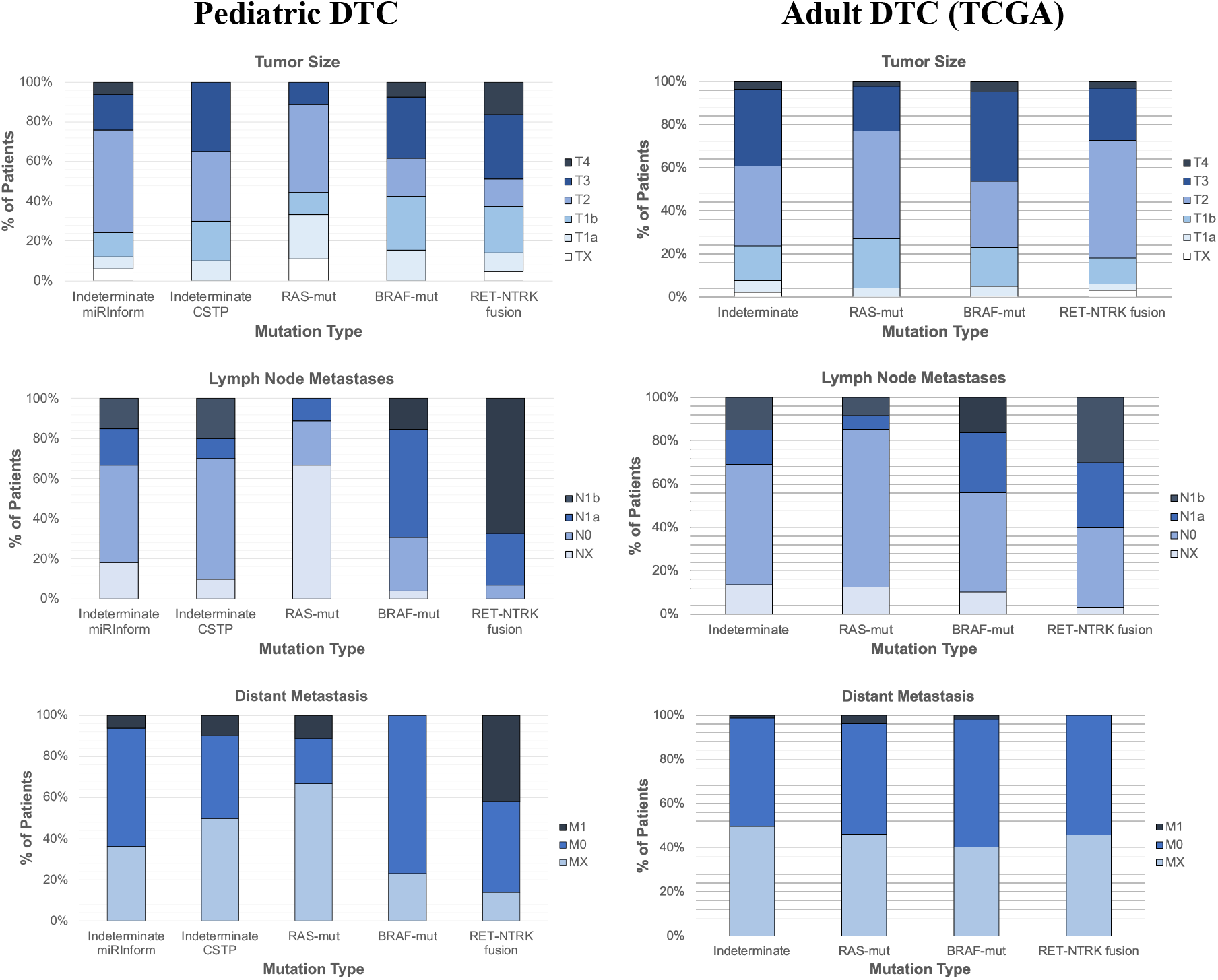
TNM staging of pediatric and adult thyroid cancers according to mutational status. Tumor Size, Lymph Node Metastasis and distant metastasis classification of pediatric differentiated thyroid cancer (n=131; combined CSTP and miRInform samples) and adult differentiated thyroid cancers (n=496; TCGA) categorized by Indeterminate, *RAS*-mut, *BRAF*-mut, and *RET/NTRK* fusion variants.

*RET/NTRK* fusions were more common in our cohort of patients less than 10 years old compared to patients > 10 years old, as previously reported.^13^ Ten out of 11 (91%) patients < 10 years harbored fusion events. The prevalence gradually decreased in pediatric patients older than 10 years (27%) and into adulthood (9%). In contrast, only 1 BRAF p.V600E mutation (9%) was found among patients < 10 years old compared to 25 pediatric patients ≥ 10 years of age (20%) with further increased prevalence in adult patients with *BRAF*-mutations where 58% of PTC in adults harboring a *BRAF*-mutation (**Table 1** and **Supplementary Figure 2A**).

## DISCUSSION

The Thyroid Cancer Genome Atlas (TCGA) reported that classification of adult PTC into molecularly-defined groups, *RAS*-like and *BRAF*-like, more accurately reflects cellular signaling, cellular differentiation and clinical behavior when compared to histology alone.^1^ Our data suggests that separating pediatric PTC with *BRAF p.V600E* mutations from PTC with *RET/NTRK* fusions more closely aligns with clinicopathological features with *BRAF* positive PTC less common with decreased age and *RET/NTRK* fusion positive PTC more commonly associated with lateral neck (N1b) and distant (M1, pulmonary) metastasis. Both *BRAF* and *RET/NTRK* fusions were identified in cPTC, however, even within the same pathological variant, the molecular driver more accurately predicted metastatic behavior (**Supplementary Table 3**). This observation is in keeping with previous reports in adults^1,2,11^ and pediatrics^12,13^ that reclassification of thyroid cancers into genetic and molecular subtypes provides an opportunity for better informed clinical management compared to pathological classification alone. RAS-mutants showed similar predictability, associated with reduced metastatic behavior in our pediatric cohort compared to published data in adults, however, the low incidence of *RAS*-mutants in our cohort prevented clinically relevant statistical analysis.

There are differences in the composition of genetic variants in the pediatric population when compared to the adult. The incidence of a *BRAF p.V600E* mutation in pediatric patients with PTC is lower and there is a lower risk for *BRAF*-associated widely invasive disease with decreased radioactive iodine avidity.^5,12,13^ In addition, coexisting mutations in the *TERT* promoter, *TP53* and genes encoding effectors of the PI3K pathway (*PIK3CA*) are frequent in *BRAF*-mut (~10%) and RET/NTRK (~2.5%) advanced adult thyroid tumors.^1^ In our cohort of pediatric tumors screened by the CSTP panel (n=66), we did not find any coexisting mutations in *BRAF*-mut or *RET/NTRK* tumors. This suggests that with increasing age other age-related host factors may result in the accumulation of additional genetic alterations that negatively influence differentiation, response to therapy, and, subsequently, disease-specific morbidity and mortality supporting separation of how oncogenic landscape data is interpreted and incorporated into clinical practice for pediatric versus adult patients with DTC.

The availability of the CSTP has provided us with a wider lens in which to view pediatric thyroid carcinoma. The incorporation of a comprehensive cancer gene panel lowered the number of samples without an identifiable alteration and led to discovery of several findings that warrant further investigation. We demonstrate that 50% of tumors with distal metastasis had indeterminate drivers utilizing the miRInform panel. By contrast, only 30% of tumors characterized with the more comprehensive CSTP panel had indeterminate drivers and within these 20 samples, 12 harbored mutations that likely have an important role in thyroid tumorigenesis. These 12 cases were included in the indeterminate subgroup due to their relatively low prevalence and uncertain molecular category. These included fusions *PRKD2-BRAF, TFG-MET* (*n=2*) and *TG-FGFR1* and mutations *TSHR p.M453T* and *p.D633Y* and *BRAF p.T599del*), all previously reported in thyroid tumors with exception of the novel *PRKD2-BRAF* fusion (**Supplementary Table 2**).^1,17–19^ Future studies are underway to more clearly define the influence of these genetic alterations on altered signaling pathways and thyroid cell differentiation.

The observation of a higher incidence of *RET/NTRK* fusions as well as their association with more metastatic behavior compared to *BRAF* in our pediatric cohort emphasizes the importance of expanding our knowledge of the pediatric DTC molecular landscape. In the adult-based TCGA, analysis revealed a fairly quiet adult PTC genome allowing for a more precise evaluation of the effects of the genetic drivers on signaling pathway activation and differentiation. In the TCGA, a 71-gene signature generated by comparison of *BRAF*-mut and *RAS*-mut tumors was used to construct a *BRAF*^V600E^-*RAS* score (BRS) that separated tumors based on MAPK pathway output and clinical behavior. In the TCGA analysis, *NTRK1/3* fusions were largely neutral and virtually all *RET* fusions were only weakly *BRAF*-like.^1^ The *RET*-fusion subgroup in these adult tumors also exhibited an intermediate Thyroid Differentiation Score (TDS; 16-gene signature including thyroid specific genes such as *SLC5A5, TG, TPO, PAX8, TSHR* and others), lower than the well differentiated *RAS*-mut (*H/K/NRAS* and *PAX8/PPARG*) tumors and higher than the *BRAF*-mut (*BRAF p.V600E;* **Supplementary Figure 2**). Considering the high prevalence of *RET/NTRK* fusions in pediatric DTC, and their association with more metastatic behavior, it will be crucial to generate the transcriptional signatures of *RET/NTRK* and *BRAF*-mutant subgroups in the pediatric population to understand the differential impact of these alterations on signaling pathways, differentiation and clinical outcomes.

There are several limitations to this study, supporting the need for multi-center, prospective studies to expand our knowledge on the potential use of genetic and molecular analysis for stratification of therapy. The sample size is relatively small at 131 samples, especially as the analysis was divided between the miRInform and CSTP samples at 65 and 66 samples, respectively. It is also worth repeating that *RET/NTRK* prevalence in this study may be underestimated as the miRInform panel did not include analysis for *NTRK1, NTRK3, BRAF* and *ALK* fusions as well as expanded *RET* fusion isoforms. Unfortunately, as previously stated, the samples initially analyzed by the miRInform panel were not available for reanalysis using the CSTP. Last, our conclusions are only based on oncogenic driver alterations. Additional studies are ongoing to define the differentiation-score based on multiplatform analysis including RNA and microRNA expression.

An important strength of our study is that many of these oncogenic events that are more common in pediatrics compared to adults with DTC can now be targeted with FDA-approved agents. Of particular note is the identification of *RET* and *NTRK* fusions and positive association of these events with persistent disease and metastatic spread. Larotrectinib (NTRK inhibitor) and selpercatinib (RET inhibitor) have shown dramatic efficacy in clinical trials in both solid tumors and hematologic malignancies, including a limited number of thyroid tumors harboring *NTRK* and *RET* fusions included in these studies. Significantly, these studies have shown durable responses in a multitude of patients with limited adverse events.^20,21^ Interestingly, larotrectinib was shown to induce partial response and restore iodine uptake in one case of RAI refractory adult thyroid cancer harboring the *EML4-NTRK3* fusion^22^ opening the way for similar therapies in pediatric tumors where these alterations are frequent and often associated with poor prognosis.

## CONCLUSIONS

The combined dataset used in this study represents an evolutionary change in the information gained from genetic and molecular analysis over the span of a few short years. Based on our data, categorizing pediatric DTC into *RAS*-mutant, *BRAF*-mutant, and *RET/NTRK* fusion variants more accurately separates the higher risk of invasive behavior for *RET/NTRK* fusion driven PTC compared to PTC harboring *BRAF p.V600E* mutations. *RET/NTRK* fusion tumors metastasize to lateral neck lymph nodes at a significantly higher frequency than *BRAF p.V600E* PTC. Further, we did not observe distant metastasis in any patients with BRAF *p.V600E* mutations. These findings support the incorporation of somatic cancer gene analysis to improve the diagnostic accuracy for FNA as well as the potential utility to incorporate oncogenic data to stratify the surgical approach and to identify tumors that may benefit from oncogene-specific systemic therapies. Additional studies are underway to define the differences in differentiation score between BRAF *p.V600E* and RET/NTRK fusion PTC, differences between pediatric and adults PTC with the same oncogenic alterations, as well as to confirm the tumorigenic potential of the novel-alterations identified using the comprehensive CSTP panel.

## Abbreviations

CHOP: The Children’s Hospital of Philadelphia
CCFP: CHOP Cancer Fusion Panel
CNA: Copy Number Alteration
CSTP: CHOP Solid Tumor Panel
cmvPTC: Cribriform-Morular Variant Papillary Thyroid Cancer
cPTC: Classic Papillary Thyroid Cancer
DTC: Differentiated Thyroid Cancer
dsvPTC: Diffuse Sclerosing Variant Papillary Thyroid Cancer
ETE: Extra Thyroidal Extension
FNA: Fine Needle Aspiration
FTC: Follicular Thyroid Cancer
fvPTC: Follicular Variant Papillary Thyroid Cancer
fvPTMC: Follicular Variant Papillary Thyroid Microcarcinoma
LNM: Lymph Node Metastasis
NGS: Next Generation Sequencing
PCR: Polymerase Chain Reaction
SNV: Single-Nucleotide Variation
svPTC: Solid Variant Papillary Thyroid Cancer
tcvPTC: Tall Cell Variant Papillary Thyroid Cancer
WLPTC: Warthin-like Papillary Thyroid Cancer

## DECLARATIONS

### Ethics approval and consent to participate

The study was approved by CHOP’s Institutional Review Board.

### Consent for publication

### Availability of data and material

### Corresponding Author

Andrew J Bauer, MD

The Thyroid Center

Division of Endocrinology and Diabetes, The Children’s Hospital of Philadelphia,

3500 Civic Center Boulevard, Buerger Center 12-149, Philadelphia, PA 19104

Tel: 215-590-5129

