## Supplementary Material for "Fusion-oncogenes are associated with increased metastatic capacity and persistent disease in pediatric thyroid cancers"

**SUPPLEMENTARY TABLES**

**Supplementary Table 1.** List of cancer genes sequenced by the CHOP Solid Tumor Panel (CSTP; Targeted exome sequencing analysis), CHOP Cancer Fusion Panel (CCFP; RNA Fusion oncogene analysis) and Asuragen’s first-generation thyroid test, miRInform® Thyroid.

| # | **CSTP (DNA)** | **CCFP (RNA)** | **miRInform Thyroid Test (Asuragen)** |
| --- | --- | --- | --- |
| 1 | *ABL1* | *ABL1* | *BRAF* c.1799T>A, p.V600E |
| 2 | *ACVR1* | *ABL2* | *HRAS* c.35G>T, p.G12V |
| 3 | *AKT1* | *AKT3* | *HRAS* c.181C>A, p.Q61K |
| 4 | *AKT2* | *ALK* | *HRAS* c.182A>G, p.Q61R |
| 5 | *AKT3* | *ARHGAP26* | *KRAS* c.35G>C, p.G12A |
| 6 | *ALK* | *AXL* | *KRAS* c.34G>T, p.G12C |
| 7 | *APC* | *BCL2* | *KRAS* c.35G>A, p.G12D |
| 8 | *AR* | *BCL6* | *KRAS* c.34G>C, p.G12R |
| 9 | *ARAF* | *BCR* | *KRAS* c.34G>A, p.G12S |
| 10 | *ARID1A* | *BRAF* | *KRAS* c.35G>T, p.G12V |
| 11 | *ARID1B* | *BRD3* | *KRAS* c.38G>A, p.G13D |
| 12 | *ARID2* | *BRD4* | *NRAS* c.181C>A, p.Q61K |
| 13 | *ASXL1* | *CAMTA1* | *NRAS* c.182A>T, p.Q61L |
| 14 | *ATM* | *CBFB* | *NRAS* c.182A>G, p.Q61R |
| 15 | *ATR* | *CCNB3* | *PAX8-PPARG* |
| 16 | *ATRX* | *CCND1* | *RET-PTC1* (*CCDC6-RET*) |
| 17 | *AURKA* | *CIC* | *RET-PTC3* (*NCOA4-RET*) |
| 18 | *AURKB* | *CRFL2* |  |
| 19 | *AXIN1* | *CSF1R* |  |
| 20 | *AXL* | *DNAJB1* |  |
| 21 | *B2M* | *DUSP22* |  |
| 22 | *BAP1* | *EGFR* |  |
| 23 | *BARD1* | *EPC1* |  |
| 24 | *BCL2* | *EPOR* |  |
| 25 | *BCL6* | *ERG* |  |
| 26 | *BCOR* | *ESR1* |  |
| 27 | *BCORL1* | *ESRRA* |  |
| 28 | *BLM* | *ETV1* |  |
| 29 | *BRAF* | *ETV4* |  |
| 30 | *BRCA1* | *ETV5* |  |
| 31 | *BCRA2* | *ETV6* |  |
| 32 | *BRD4* | *EWSR1* |  |
| 33 | *BRIP1* | *FGFR1* |  |
| 34 | *CARD11* | *FGFR2* |  |
| 35 | *CBFB* | *FGFR3* |  |
| 36 | *CBL* | *FGR* |  |
| 37 | *CCND1* | *FOXO1* |  |
| 38 | *CCND2* | *FUS* |  |
| 39 | *CCND3* | *GLI1* |  |
| 40 | *CCNE1* | *GLIS2* |  |
| 41 | *CD274* | *HMGA2* |  |
| 42 | *CD79B* | *IL2RB* |  |
| 43 | *CDC73* | *IL3* |  |
| 44 | *CDH1* | *INSR* |  |
| 45 | *CDK12* | *JAK2* |  |
| 46 | *CDK4* | *JAZF1* |  |
| 47 | *CDK6* | *KMT2A* |  |
| 48 | *CDK8* | *MALT1* |  |
| 49 | *CDKN1B* | *MAML2* |  |
| 50 | *CDKN2A* | *MAST1* |  |
| 51 | *CDKN2B* | *MAST2* |  |
| 52 | *CDKN2C* | *MEAF6* |  |
| 53 | *CHEK1* | *MECOM* |  |
| 54 | *CHEK2* | *MET* |  |
| 55 | *CIC* | *MKL1* |  |
| 56 | *CREBBP* | *MKL2* |  |
| 57 | *CRKL* | *MSMB* |  |
| 58 | *CRLF2* | *MUSK* |  |
| 59 | *CSF1R* | *MYB* |  |
| 60 | *CTCF* | *MYC* |  |
| 61 | *CTNNB1* | *NCOA2* |  |
| 62 | *DAXX* | *NOTCH1* |  |
| 63 | *DDR2* | *NOTCH2* |  |
| 64 | *DICER1* | *NRG1* |  |
| 65 | *DNMT3A* | *NTRK1* |  |
| 66 | *DOT1L* | *NTRK2* |  |
| 67 | *EED* | *NTRK3* |  |
| 68 | *EGFR* | *NUMBL* |  |
| 69 | *EP300* | *NUP214* |  |
| 70 | *EPHA3* | *NUP98* |  |
| 71 | *EPHA5* | *NUT* |  |
| 72 | *EPHB1* | *PAX5* |  |
| 73 | *ERBB2* | *PDGFB* |  |
| 74 | *ERBB3* | *PDGFRA* |  |
| 75 | *ERBB4* | *PDGFRB* |  |
| 76 | *ERG* | *PICALM* |  |
| 77 | *ESR1* | *PIK3CA* |  |
| 78 | *ETV6* | *PKN1* |  |
| 79 | *EZH2* | *PLAG1* |  |
| 80 | *FAM46C* | *PPARG* |  |
| 81 | *FANCA* | *PRKACA* |  |
| 82 | *FANCC* | *PRKCA* |  |
| 83 | *FBXW7* | *PRKCB* |  |
| 84 | *FGF19* | *PTK2B* |  |
| 85 | *FGF3* | *RAF1* |  |
| 86 | *FGF4* | *RARA* |  |
| 87 | *FGFR1* | *RBM15* |  |
| 88 | *FGFR2* | *RELA* |  |
| 89 | *FGFR3* | *RET* |  |
| 90 | *FGFR4* | *ROS1* |  |
| 91 | *FLCN* | *RSPO2* |  |
| 92 | *FLT1* | *RSPO3* |  |
| 93 | *FLT3* | *RUNX1* |  |
| 94 | *FLT4* | *RUNX1T1* |  |
| 95 | *FOXL2* | *SS18* |  |
| 96 | *FOXP1* | *STAT6* |  |
| 97 | *FUBP1* | *TAF15* |  |
| 98 | *GATA1* | *TAL1* |  |
| 99 | *GATA2* | *TCF12* |  |
| 100 | *GATA3* | *TCF3* |  |
| 101 | *GNA11* | *TERT* |  |
| 102 | *GNAQ* | *TFE3* |  |
| 103 | *GNAS* | *TFEB* |  |
| 104 | *GRIN2A* | *TFG* |  |
| 105 | *GSK3B* | *THADA* |  |
| 106 | *H3F3A* | *TMPRSS2* |  |
| 107 | *HGF* | *TSLP* |  |
| 108 | *HIST1H1C* | *TYK2* |  |
| 109 | *HIST1H3B* | *USP6* |  |
| 110 | *HNF1A* | *YWHAE* |  |
| 111 | *HRAS* |  |  |
| 112 | *IDH1* |  |  |
| 113 | *IDH2* |  |  |
| 114 | *IGF1R* |  |  |
| 115 | *IKBKE* |  |  |
| 116 | *IKZF1* |  |  |
| 117 | *IL7R* |  |  |
| 118 | *INPP4B* |  |  |
| 119 | *IRF4* |  |  |
| 120 | *IRS2* |  |  |
| 121 | *JAK1* |  |  |
| 122 | *JAK2* |  |  |
| 123 | *JAK3* |  |  |
| 124 | *JMJD1C* |  |  |
| 125 | *JUN* |  |  |
| 126 | *KDM5A* |  |  |
| 127 | *KDM5C* |  |  |
| 128 | *KDM6A* |  |  |
| 129 | *KDR* |  |  |
| 130 | *KEAP1* |  |  |
| 131 | *KIT* |  |  |
| 132 | *KMT2A* |  |  |
| 133 | *KMT2C* |  |  |
| 134 | *KRAS* |  |  |
| 135 | *MAP2K1* |  |  |
| 136 | *MAP2K2* |  |  |
| 137 | *MAP2K4* |  |  |
| 138 | *MAP3K1* |  |  |
| 139 | *MAPK1* |  |  |
| 140 | *MCL1* |  |  |
| 141 | *MDM2* |  |  |
| 142 | *MDM4* |  |  |
| 143 | *MED12* |  |  |
| 144 | *MEF2B* |  |  |
| 145 | *MEN1* |  |  |
| 146 | *MET* |  |  |
| 147 | *MITF* |  |  |
| 148 | *MLH1* |  |  |
| 149 | *MPL* |  |  |
| 150 | *MRE11A* |  |  |
| 151 | *MSH2* |  |  |
| 152 | *MSH6* |  |  |
| 153 | *MTOR* |  |  |
| 154 | *MUTYH* |  |  |
| 155 | *MYB* |  |  |
| 156 | *MYC* |  |  |
| 157 | *MYCN* |  |  |
| 158 | *MYD88* |  |  |
| 159 | *MYOD1* |  |  |
| 160 | *NF1* |  |  |
| 161 | *NF2* |  |  |
| 162 | *NFE2L2* |  |  |
| 163 | *NKX2-1* |  |  |
| 164 | *NOTCH1* |  |  |
| 165 | *NOTCH2* |  |  |
| 166 | *NPM1* |  |  |
| 167 | *NRAS* |  |  |
| 168 | *NTRK1* |  |  |
| 169 | *NTRK2* |  |  |
| 170 | *NTRK3* |  |  |
| 171 | *PALB2* |  |  |
| 172 | *PAX5* |  |  |
| 173 | *PBRM1* |  |  |
| 174 | *PDCD1* |  |  |
| 175 | *PDGFRA* |  |  |
| 176 | *PDGFRB* |  |  |
| 177 | *PHOX2B* |  |  |
| 178 | *PIK3CA* |  |  |
| 179 | *PIK3CG* |  |  |
| 180 | *PIK3R1* |  |  |
| 181 | *PIK3R2* |  |  |
| 182 | *PIM1* |  |  |
| 183 | *PPM1D* |  |  |
| 184 | *PPP2R1A* |  |  |
| 185 | *PRDM1* |  |  |
| 186 | *PRKAR1A* |  |  |
| 187 | *PTCH1* |  |  |
| 188 | *PTEN* |  |  |
| 189 | *PTPN11* |  |  |
| 190 | *RAD50* |  |  |
| 191 | *RAD51* |  |  |
| 192 | *RAF1* |  |  |
| 193 | *RARA* |  |  |
| 194 | *RB1* |  |  |
| 195 | *RET* |  |  |
| 196 | *RHOA* |  |  |
| 197 | *RICTOR* |  |  |
| 198 | *RNF43* |  |  |
| 199 | *ROS1* |  |  |
| 200 | *RPTOR* |  |  |
| 201 | *RUNX1* |  |  |
| 202 | *SDHA* |  |  |
| 203 | *SDHB* |  |  |
| 204 | *SDHC* |  |  |
| 205 | *SDHD* |  |  |
| 206 | *SETD2* |  |  |
| 207 | *SF3B1* |  |  |
| 208 | *SMAD2* |  |  |
| 209 | *SMAD4* |  |  |
| 210 | *SMARCA4* |  |  |
| 211 | *SMARCB1* |  |  |
| 212 | *SMO* |  |  |
| 213 | *SOCS1* |  |  |
| 214 | *SOX2* |  |  |
| 215 | *SPEN* |  |  |
| 216 | *SPOP* |  |  |
| 217 | *SRC* |  |  |
| 218 | *STAG2* |  |  |
| 219 | *STK11* |  |  |
| 220 | *SUFU* |  |  |
| 221 | *SUZ12* |  |  |
| 222 | *TERT* |  |  |
| 223 | *TET2* |  |  |
| 224 | *TGFBR2* |  |  |
| 225 | *TNFAIP3* |  |  |
| 226 | *TNFRSF14* |  |  |
| 227 | *TOP1* |  |  |
| 228 | *TP53* |  |  |
| 229 | *TP63* |  |  |
| 230 | *TSC1* |  |  |
| 231 | *TSC2* |  |  |
| 232 | *TSHR* |  |  |
| 233 | *U2AF1* |  |  |
| 234 | *VHL* |  |  |
| 235 | *WHSC1* |  |  |
| 236 | *WT1* |  |  |
| 237 | *AMER1* |  |  |
| 238 | *XPO1* |  |  |

**Supplementary Table 2.** List of driver genetic alterations found in 131 pediatric thyroid cancers using miRInform Thyroid Test (n=65) and CSTP/CCFP panels (n=66).

| **ID** | **Sex** | **Age** | **Histology** | | **Panel** | | ***Gene*** | | **Alteration** | | **Molecular category** | |
| --- | --- | --- | --- | --- | --- | --- | --- | --- | --- | --- | --- | --- |
| 1 | F | ≥10 | | FTC | | miR*Inform* | |  | |  | | Indeterminate |
| 2 | F | ≥10 | | FTC | | miR*Inform* | |  | |  | | Indeterminate |
| 3 | F | ≥10 | | FTC | | miR*Inform* | |  | |  | | Indeterminate |
| 4 | F | ≥10 | | FTC | | miR*Inform* | | *KRAS* | | p.G12V | | RAS-mut |
| 5 | M | ≥10 | | FTC | | miR*Inform* | |  | |  | | Indeterminate |
| 6 | F | ≥10 | | cPTC | | miR*Inform* | | *BRAF* | | p.V600E | | BRAF-mut |
| 7 | F | ≥10 | | cPTC | | miR*Inform* | | *BRAF* | | p.V600E | | BRAF-mut |
| 8 | F | ≥10 | | cPTC/fvPTC | | miR*Inform* | | *BRAF* | | p.V600E | | BRAF-mut |
| 9 | F | ≥10 | | cPTC | | miR*Inform* | | *BRAF* | | p.V600E | | BRAF-mut |
| 10 | F | ≥10 | | cPTC | | miR*Inform* | | *BRAF* | | p.V600E | | BRAF-mut |
| 11 | F | ≥10 | | cPTC | | miR*Inform* | | *BRAF* | | p.V600E | | BRAF-mut |
| 12 | F | ≥10 | | cPTC/fvPTC | | miR*Inform* | |  | |  | | Indeterminate |
| 13 | F | ≥10 | | cPTC | | miR*Inform* | |  | |  | | Indeterminate |
| 14 | F | ≥10 | | cPTC | | miR*Inform* | | *RET* | | *CCDC6-RET* | | RET/NTRK fusion |
| 15 | M | ≥10 | | cPTC | | miR*Inform* | | *BRAF* | | p.V600E | | BRAF-mut |
| 16 | M | ≥10 | | fvPTC | | miR*Inform* | |  | |  | | Indeterminate |
| 17 | F | ≥10 | | cPTC | | miR*Inform* | | *BRAF* | | p.V600E | | BRAF-mut |
| 18 | F | ≥10 | | fvPTC | | miR*Inform* | |  | |  | | Indeterminate |
| 19 | M | ≥10 | | fvPTC | | miR*Inform* | |  | |  | | Indeterminate |
| 20 | F | ≥10 | | fvPTC | | miR*Inform* | | *NRAS* | | p.Q61R | | RAS-mut |
| 43 | F | ≥10 | | fvPTC | | miR*Inform* | |  | |  | | Indeterminate |
| 44 | M | ≥10 | | cPTC | | miR*Inform* | | *RET* | | *CCDC6-RET* | | RET/NTRK fusion |
| 45 | F | ≥10 | | cPTC/fvPTC | | miR*Inform* | | *BRAF* | | p.V600E | | BRAF-mut |
| 46 | F | ≥10 | | cPTC | | miR*Inform* | | *RET* | | *CCDC6-RET* | | RET/NTRK fusion |
| 47 | F | ≥10 | | dsvPTC | | miR*Inform* | | *RET* | | *CCDC6-RET* | | RET/NTRK fusion |
| 48 | F | ≥10 | | dsvPTC | | miR*Inform* | | *RET* | | *CCDC6-RET* | | RET/NTRK fusion |
| 49 | F | ≥10 | | dsvPTC | | miR*Inform* | | *RET* | | *CCDC6-RET* | | RET/NTRK fusion |
| 50 | M | ≥10 | | Other: cPTC/fvPTC/svPTC | | miR*Inform* | | *PAX8/PPARg* | | *PAX8/PPARg* | | RAS-mut |
| 51 | F | ≥10 | | fvPTC | | miR*Inform* | |  | |  | | Indeterminate |
| 58 | M | ≥10 | | cPTC | | miR*Inform* | |  | |  | | Indeterminate |
| 59 | F | ≥10 | | cPTC | | miR*Inform* | |  | |  | | Indeterminate |
| 61 | M | <10 | | cPTC | | miR*Inform* | | *BRAF* | | p.V600E | | BRAF-mut |
| 62 | F | ≥10 | | cPTC | | miR*Inform* | |  | |  | | Indeterminate |
| 63 | F | <10 | | cPTC | | miR*Inform* | | *RET* | | *CCDC6-RET* | | RET/NTRK fusion |
| 65 | F | ≥10 | | cPTC | | miR*Inform* | |  | |  | | Indeterminate |
| 66 | F | ≥10 | | dsvPTC | | miR*Inform* | |  | |  | | Indeterminate |
| 67 | M | ≥10 | | dsvPTC | | miR*Inform* | |  | |  | | Indeterminate |
| 68 | F | ≥10 | | dsvPTC | | miR*Inform* | |  | |  | | Indeterminate |
| 69 | F | <10 | | dsvPTC | | miR*Inform* | | *RET* | | *CCDC6-RET* | | RET/NTRK fusion |
| 71 | F | ≥10 | | fvPTC | | miR*Inform* | |  | |  | | Indeterminate |
| 72 | F | ≥10 | | FTC | | miR*Inform* | | *HRAS* | | p.Q61R | | RAS-mut |
| 73 | M | ≥10 | | fvPTC | | miR*Inform* | | *PAX8/PPARg* | | *PAX8/PPARg* | | RAS-mut |
| 74 | M | ≥10 | | fvPTC | | miR*Inform* | | *NRAS* | | p.Q61R | | RAS-mut |
| 75 | F | ≥10 | | fvPTC | | miR*Inform* | |  | |  | | Indeterminate |
| 76 | F | ≥10 | | fvPTC | | miR*Inform* | |  | |  | | Indeterminate |
| 77 | F | ≥10 | | fvPTC | | miR*Inform* | |  | |  | | Indeterminate |
| 78 | F | ≥10 | | fvPTC | | miR*Inform* | |  | |  | | Indeterminate |
| 79 | F | ≥10 | | fvPTC | | miR*Inform* | | *NRAS* | | p.Q61R | | RAS-mut |
| 80 | F | ≥10 | | fvPTC | | miR*Inform* | | *RET* | | *CCDC6-RET* | | RET/NTRK fusion |
| 81 | F | ≥10 | | cPTC | | miR*Inform* | | *RET* | | *CCDC6-RET* | | RET/NTRK fusion |
| 82 | F | ≥10 | | Other: cPTC/fvPTC/svPTC | | miR*Inform* | |  | |  | | Indeterminate |
| 83 | M | ≥10 | | fvPTC | | miR*Inform* | |  | |  | | Indeterminate |
| 84 | F | ≥10 | | fvPTC | | miR*Inform* | |  | |  | | Indeterminate |
| 85 | F | ≥10 | | fvPTC | | miR*Inform* | |  | |  | | Indeterminate |
| 100 | M | <10 | | Other: cPTC/fvPTC/svPTC | | miR*Inform* | | *RET* | | *NCOA4-RET* | | RET/NTRK fusion |
| 101 | F | ≥10 | | cPTC | | miR*Inform* | | *BRAF* | | p.V600E | | BRAF-mut |
| 103 | M | ≥10 | | fvPTC | | miR*Inform* | |  | |  | | Indeterminate |
| 104 | F | ≥10 | | cPTC | | miR*Inform* | |  | |  | | Indeterminate |
| 107 | M | ≥10 | | Other: fvPTC/tcvPTC | | miR*Inform* | |  | |  | | Indeterminate |
| 108 | M | ≥10 | | fvPTC | | miR*Inform* | | *NRAS* | | p.Q61R | | RAS-mut |
| 110 | F | ≥10 | | cPTC | | miR*Inform* | |  | |  | | Indeterminate |
| 111 | F | ≥10 | | fvPTC | | miR*Inform* | |  | |  | | Indeterminate |
| 112 | F | ≥10 | | cPTC | | miR*Inform* | | *BRAF* | | p.V600E | | BRAF-mut |
| 114 | F | ≥10 | | cPTC | | miR*Inform* | | *RET* | | *NCOA4-RET* | | RET/NTRK fusion |
| 115 | M | ≥10 | | cPTC | | miR*Inform* | |  | |  | | Indeterminate |
| D01 | M | ≥10 | | dsvPTC | | CSTP | | *RET* | | *CCDC6-RET* | | RET/NTRK fusion |
| D02 | F | ≥10 | | fvPTC | | CSTP | | TSHR | | p.D633Y | | Indeterminate |
| D03 | M | <10 | | dsvPTC | | CSTP | | *NTRK1* | | *IRF2BP-NTRK1* | | RET/NTRK fusion |
| D04 | F | <10 | | fvPTC | | CSTP | | *NTRK3* | | *ETV6-NTRK3* | | RET/NTRK fusion |
| D05 | F | ≥10 | | dsvPTC | | CSTP | | *RET* | | *CCDC6-RET* | | RET/NTRK fusion |
| D06 | F | ≥10 | | Other: cmvPTC | | CSTP | | *APC* | | Homozygous deletion | | Indeterminate |
| D07 | F | ≥10 | | cPTC | | CSTP | |  | |  | | Indeterminate |
| D08 | F | ≥10 | | dsvPTC | | CSTP | | *RET* | | *CCDC6-RET* | | RET/NTRK fusion |
| D09 | F | ≥10 | | cPTC | | CSTP | | *RET* | | *PRKAR1A-RET* | | RET/NTRK fusion |
| D10 | F | ≥10 | | cPTC | | CSTP | | *NTRK3* | | *ETV6-NTRK3* | | RET/NTRK fusion |
| D11 | M | <10 | | dsvPTC | | CSTP | | *RET* | | *SPECC1L-RET* | | RET/NTRK fusion |
| D12 | F | ≥10 | | fvPTC | | CSTP | | *BRAF* | | p.V600E | | BRAF-mut |
| D13 | F | ≥10 | | cPTC/fvPTC | | CSTP | |  | |  | | Indeterminate |
| D14 | F | <10 | | cPTC | | CSTP | | *RET* | | *NCOA4-RET* | | RET/NTRK fusion |
| D15 | M | ≥10 | | Other: svPTC | | CSTP | | *RET* | | *CCDC6-RET* | | RET/NTRK fusion |
| D16 | F | ≥10 | | cPTC | | CSTP | | *BRAF* | | p.V600E | | BRAF-mut |
| D17 | F | ≥10 | | cPTC | | CSTP | | *BRAF* | | p.V600E | | BRAF-mut |
| D18 | F | ≥10 | | cPTC | | CSTP | | *BRAF* | | p.V600E | | BRAF-mut |
| D19 | F | <10 | | cPTC | | CSTP | | *NTRK1* | | *TPR-NTRK1* | | RET/NTRK fusion |
| D20 | M | ≥10 | | cPTC | | CSTP | | *RET* | | *CCDC6-RET* | | RET/NTRK fusion |
| D21 | F | ≥10 | | FTC | | CSTP | | *DICER1* | | p. G661Vfs*24,  p. E1813K | | Indeterminate |
| D22 | F | ≥10 | | cPTC | | CSTP | | *NTRK3* | | *ETV6-NTRK3* | | RET/NTRK fusion |
| D23 | M | ≥10 | | fvPTC | | CSTP | | *NTRK3* | | *ETV6-NTRK3* | | RET/NTRK fusion |
| D24 | M | ≥10 | | dsvPTC | | CSTP | | *RET* | | *CCDC6-RET* | | RET/NTRK fusion |
| D25 | F | ≥10 | | cPTC | | CSTP | | *RET* | | *CCDC6-RET* | | RET/NTRK fusion |
| D26 | F | ≥10 | | cPTC | | CSTP | | *BRAF* | | p.V600E | | BRAF-mut |
| D28 | F | ≥10 | | fvPTC | | CSTP | | *DICER1* | | p.E1705K  p.L777fs | | Indeterminate |
| D29 | F | ≥10 | | FTC | | CSTP | |  | |  | | Indeterminate |
| D30 | F | ≥10 | | cPTC | | CSTP | | *RET* | | *NCOA4-RET* | | RET/NTRK fusion |
| D31 | F | <10 | | cPTC | | CSTP | | *NTRK1* | | *SQSTM1-NTRK1* | | RET/NTRK fusion |
| D32 | F | ≥10 | | cPTC | | CSTP | | *NTRK3* | | *ETV6-NTRK3* | | RET/NTRK fusion |
| D33 | F | ≥10 | | cPTC | | CSTP | | *NRAS* | | p.Q61R | | RAS-mut |
| D34 | F | ≥10 | | cPTC/fvPTC | | CSTP | |  | |  | | Indeterminate |
| D35 | F | ≥10 | | cPTC | | CSTP | | *BRAF* | | p.V600E | | BRAF-mut |
| D36 | F | ≥10 | | cPTC | | CSTP | | *RET* | | *EML4-RET* | | RET/NTRK fusion |
| D37 | F | ≥10 | | cPTC | | CSTP | | *NTRK1* | | *TPR-NTRK1* | | RET/NTRK fusion |
| D38 | M | ≥10 | | cPTC/fvPTC | | CSTP | | *BRAF* | | p.T599del | | Indeterminate |
| D39 | M | ≥10 | | cPTC | | CSTP | | *RET* | | *NCOA4-RET* | | RET/NTRK fusion |
| D40 | F | ≥10 | | cPTC | | CSTP | | *RET* | | *CCDC6-RET* | | RET/NTRK fusion |
| D41 | F | ≥10 | | fvPTC | | CSTP | | TSHR | | p.M543T | | Indeterminate |
| D42 | M | ≥10 | | FTC | | CSTP | | *FGFR1* | | *TG-FGFR1* | | Indeterminate |
| D43 | F | ≥10 | | cPTC | | CSTP | | *MET* | | *TFG-MET* | | Indeterminate |
| D44 | F | ≥10 | | cPTC | | CSTP | | *BRAF* | | p.V600E | | BRAF-mut |
| D45 | F | ≥10 | | Other: cmvPTC | | CSTP | | *APC* | | p.S982Rfs*23 | | Indeterminate |
| D46 | F | ≥10 | | cPTC/fvPTC | | CSTP | |  | |  | | Indeterminate |
| D47 | F | ≥10 | | cPTC | | CSTP | | *BRAF* | | p.V600E | | BRAF-mut |
| D48 | F | ≥10 | | cPTC | | CSTP | | *BRAF* | | p.V600E | | BRAF-mut |
| D49 | F | ≥10 | | cPTC | | CSTP | | *BRAF* | | *PRKD2-BRAF* | | Indeterminate |
| D50 | M | ≥10 | | cPTC | | CSTP | |  | |  | | Indeterminate |
| D51 | M | ≥10 | | cPTC | | CSTP | | *BRAF* | | p.V600E | | BRAF-mut |
| D52 | F | ≥10 | | fvPTC | | CSTP | | *RET* | | *NCOA4-RET* | | RET/NTRK fusion |
| D53 | M | ≥10 | | cPTC | | CSTP | | *RET* | | *SPECC1L-RET* | | RET/NTRK fusion |
| D54 | F | ≥10 | | Other: cmvPTC | | CSTP | | *APC* | | p.R216*  c.1744-2A>G | | Indeterminate |
| D55 | F | ≥10 | | cPTC/fvPTC | | CSTP | | *RET* | | *CCDC6-RET* | | RET/NTRK fusion |
| D56 | F | ≥10 | | cPTC | | CSTP | | *BRAF* | | p.V600E | | BRAF-mut |
| D57 | F | ≥10 | | Other: cPTC/fvPTC/svPTC | | CSTP | | *RET* | | *NCOA4-RET* | | RET/NTRK fusion |
| D58 | M | <10 | | fvPTC | | CSTP | | *RET* | | *NCOA4-RET* | | RET/NTRK fusion |
| D59 | F | ≥10 | | cPTC | | CSTP | | *RET* | | *CCDC6-RET* | | RET/NTRK fusion |
| D60 | F | ≥10 | | cPTC | | CSTP | | *MET* | | *TFG-MET* | | Indeterminate |
| D61 | M | ≥10 | | Other: fvPTC/svPTC | | CSTP | | *RET* | | *CCDC6-RET* | | RET/NTRK fusion |
| D62 | F | ≥10 | | Other: WLPTC | | CSTP | | *BRAF* | | p.V600E | | BRAF-mut |
| D63 | F | ≥10 | | cPTC | | CSTP | |  | |  | | Indeterminate |
| D64 | M | ≥10 | | cPTC | | CSTP | |  | |  | | Indeterminate |
| D65 | M | ≥10 | | cPTC | | CSTP | | *BRAF* | | p.V600E | | BRAF-mut |
| D66 | F | ≥10 | | dsvPTC | | CSTP | | *RET* | | *TRIM24-RET* | | RET/NTRK fusion |
| D67 | F | ≥10 | | cPTC | | CSTP | | *BRAF* | | p.V600E | | BRAF-mut |

* Highlighted cases were classified as indeterminate due to the low prevalence and uncertain molecular category

of the driver alteration.

**Supplementary Table 3.** Association between *BRAF*-mut and *RET/NTRK* fusion subgroups and remission, lymph node metastasis (N) and distant metastasis (M) in classic variant PTC (cPTC).

|  |  | *BRAF*-mut | *RET/NTRK* | p-value, |
| --- | --- | --- | --- | --- |
|  |  |  |  | *BRAF* vs *RET/NTRK* |
| Remission | Yes | 17 (94.4%) | 10 (66.7%) | **3.90E-02** |
|  | No | 1 (5.6%) | 5 (33.3%) |  |
| N | 0 | 6 (27.3%) | 1 (4.5%) | **1.04E-02** |
|  | 1a | 12 (54.5%) | 8 (36.4%) |  |
|  | 1b | 4 (18.2%) | 13 (59.1%) |  |
| M | 0 | 17 (77.3%) | 11 (50.0%) | **7.50E-03** |
|  | 1 | 0 (0%) | 8 (36.4%) |  |
|  | X | 5 (22.7%) | 3 (13.6%) |  |

**Age**

**Group:**

**Supplementary Figure 1.** Diagnostic age of thyroid cancer patients. **A.** Median age of the different molecular subgroups (*BRAF*-mut, *RET/NTRK* fusion, *RAS*-mut and Indeterminate) in pediatric and adult thyroid cancers (TCGA). **B.** Distribution of *BRAF* mutations and *RET/NTRK* fusions across different age quartiles in adult thyroid cancer TCGA. Younger adult patients (21-36 years old) harbor 50% of all *RET/NTRK* fusions whereas *BRAF* mutations are similarly distributed between different age groups.

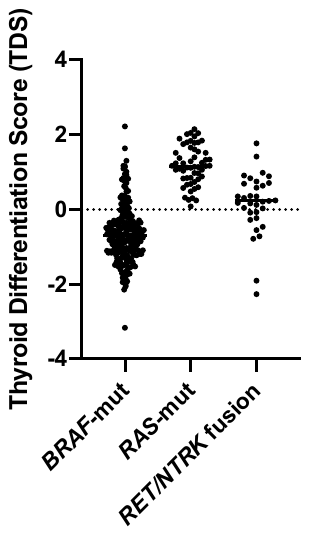

P<0.0001

Kruskal-Wallis Test

**Supplementary Figure 2.** Thyroid Differentiation Score (TDS) of different molecular subgroups (*BRAF*-mut, *RET/NTRK* fusion, *RAS*-mut) in adult thyroid cancers (TCGA).
